# Quantitative Attributions with Counterfactuals

**DOI:** 10.1101/2024.11.26.625505

**Authors:** Diane-Yayra Adjavon, Nils Eckstein, Alexander S. Bates, Gregory S.X.E. Jefferis, Jan Funke

**Affiliations:** HHMI Janelia; Harvard Medical School; MRC LMB, Cambridge

## Abstract

We address the problem of explaining the decision process of deep neural network classifiers on images, which is of particular importance in biomedical datasets where class-relevant differences are not always obvious to a human observer. Our proposed solution, termed quantitative attribution with counterfactuals (QuAC), generates visual explanations that highlight class-relevant differences by attributing the classifier decision to changes of visual features in small parts of an image. To that end, we train a separate network to generate counterfactual images (*i*.*e*., to translate images between different classes). We then find the most important differences using novel discriminative attribution methods. Crucially, QuAC allows scoring of the attribution and thus provides a measure to quantify and compare the fidelity of a visual explanation. We demonstrate the suitability and limitations of QuAC on two datasets: (1) a synthetic dataset with known class differences, representing different levels of protein aggregation in cells and (2) an electron microscopy dataset of *D. melanogaster* synapses with different neurotransmitters, where QuAC reveals so far unknown visual differences. We further discuss how QuAC can be used to interrogate mispredictions to shed light on unexpected inter-class similarities and intra-class differences.

## 1 Introduction

As more and more image analysis solutions in the life sciences rely on artificial neural networks, there is an increased need for reliable and quantifiable explainability methods to understand how those networks make decisions. Importantly, these explainability methods should provide testable hypotheses about the visual features used by a network. Knowing which features a network uses is important to build confidence by confirming that a network focuses on relevant features instead of dataset confounders. Furthermore, revealing the features involved in a decision process can lead to new insights about phenotypic differences.

Here, we address the problem of revealing which visual features a neural network uses to classify microscopy images. Our proposed solution (Quantitative Attributions with Counterfactuals, QuAC), finds classifier-relevant *differences* between pairs of classes by converting microscopy images from one class to another. The result of the conversion is a counterfactual example that differs only in the smallest possible region compared to the original image, while still being classified differently than the original image (see Fig. 1a). Crucially, our method allows us to quantify the importance of the modified region with respect to the classifier’s prediction and thus to find the most informative differences between classes.

**Figure 1.**
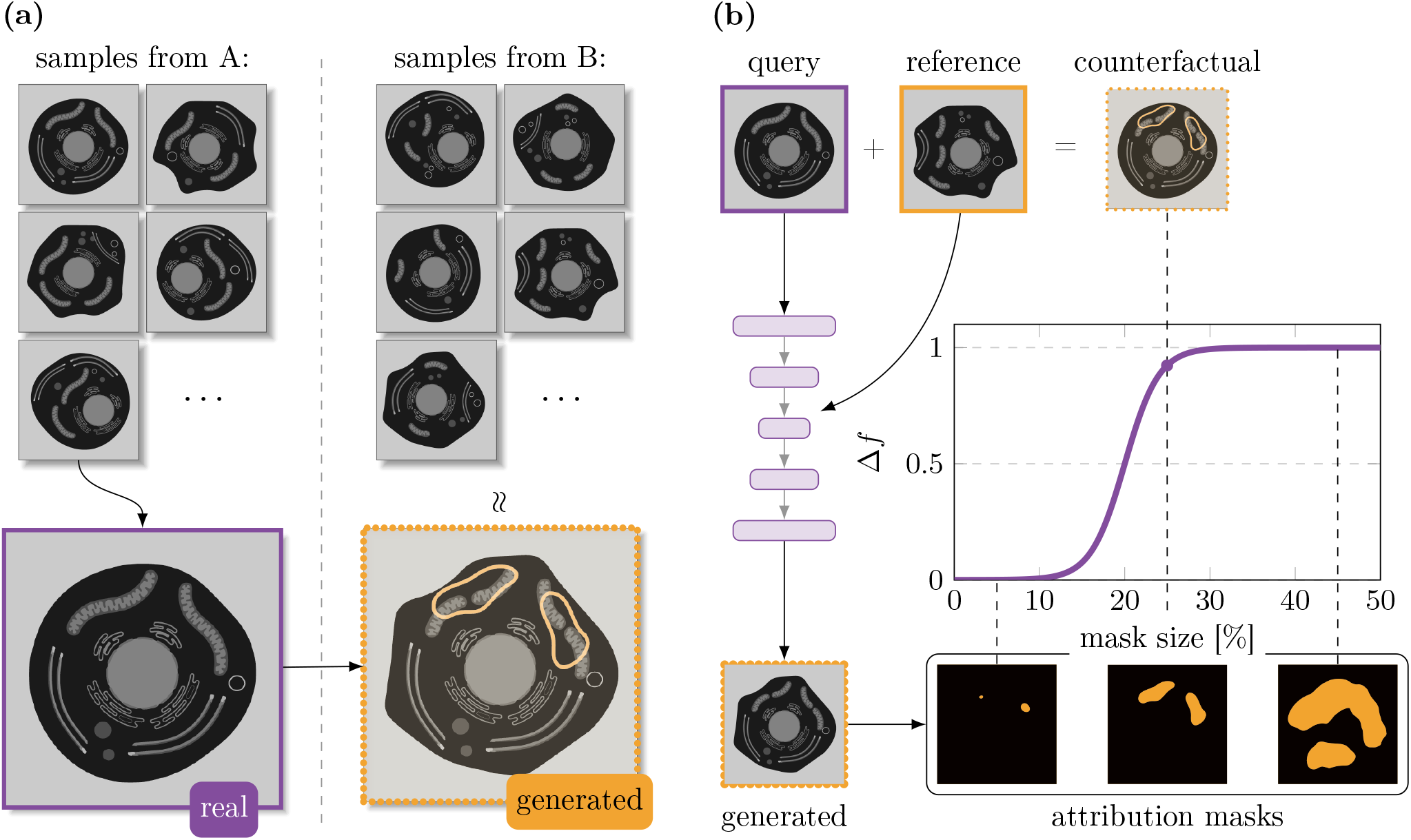
Overview of QuAC. In panel **(a)** we show a schematic example how QuAC works. The data set consists of different classes, here just two classes A and B, which can be distinguished by a classifier. Importantly, there is as much intra-class heterogeneity as inter-class heterogeneity of features; as such the class-relevant features are not evident. QuAC solves this problem by converting a sample from class A into a sample from class B, and then highlighting the difference. **(b)** QuAC gives a quantitative evaluation of how good our explanation of the classifier’s decision is. From the input of class A, the neural network suggest a generated image which is classified as class B. Using a discriminative attribution, we obtain a ranking of how important regions of the image are for the classifier’s decision. These regions are described using a binary mask, shown at the bottom - they are evaluated by how much they change the classifier’s decision. The combination of the generated image and the input image, given a binary mask that optimally describes the difference between the two images, gives us a counterfactual. The counterfactual differs from the input image only by the minimal set of regions that are necessary to change the classifier’s decision.

We consider the case where a classifier can successfully discriminate between images of different classes (possibly on subtle differences that are not obvious to human observers). To reveal which visual features the classifier uses, we first train a separate generative model to translate a *query* image of one class into a different class given by a *reference* image. The result of the translation is a generated image that is most similar to the query image, but is a representative sample of the reference image’s class (see Fig. 1b, left). We then obtain a series of attribution masks of varying size using a discriminative attribution method (Fig. 1b, bottom, and described in detail in Section 3). For each mask, we create a counterfactual image by replacing the query image with the generated image in the region of the attribution mask (Fig. 1b, top). We quantify how much these changes affect the class prediction by measuring the difference Δ*f* in the classifier’s output between the query image and each counterfactual image. When plotted against the mask size, this gives the QuAC curve (Fig. 1b, center). A QuAC curve with a sharp, early increase indicates that the classifier’s decision is sensitive to the region described by the attribution masks: we therefore use the area under the curve as the *QuAC score* for the efficiency of an attribution method. We can use the QuAC score to rank attribution methods and choose the one with the highest score. From the attribution of the chosen method, we can then identify the smallest mask that leads to the highest change in the classifier’s output Δ*f*. In Fig. 1a, this occurs at a mask size of approximately 25% of the image. The counterfactual image created with this mask is then the most informative image for the classifier’s decision, as it reveals the most important and most localized visual differences between the two classes.

We demonstrate QuAC on two datasets representing different imaging modalities and biological applications: simulated light microscopy data of cells with known phenotypical differences, and electron microscopy images of *D. melanogaster* synapses releasing different neurotransmitters. We consider classification tasks which can be difficult—or as of yet impossible—for humans to perform, and show that QuAC helps to understand the decision process of a classifier on those datasets. We argue that the explanations provided by QuAC help to verify that a classifier uses relevant features (as opposed to dataset confounders) and can even be used to discover so-far unknown phenotypical differences between classes. We show that QuAC can recapitulate all the known axes of variance used for classification of cells on simulated light microscopy (Section 2.1). QuAC provides new biological insights into the relation between structure and function of synapses, shown already using a previous verion of the method [1, 2], and expanded upon here (Section 2.2). Finally, we demonstrate in a case study on the synapse dataset how QuAC can be used to interrogate and interpret classifier mispredictions (Section 2.3).

Overall, QuAC provides detailed, realistic, and quantitative explanations that are particularly well-suited to fine-grained classification tasks—especially those which are difficult for human observers to perform. QuAC can be used to discover subtle phenotypic differences between different conditions. We hope that QuAC and other similar future methods will lead to new insights in, *e*.*g*., the relation of genotypes and phenotypes or subtle differences between diseased and healthy states of cells and tissue. In particular, we hypothesize that QuAC can be used for understanding and improving early diagnosis or early differentiation stages where the changes in the images are not obvious to the human eye. We should note that QuAC works best on images where class-relevant differences are localized. Global features (like intensity variations, if they are class relevant) would lead to overly large and potentially difficult to interpret attribution maps. A different data representation would be needed in those cases, such that relevant features are can be localized and interpreted.

## 2 Results

### 2.1 Synthetic cells

We first consider a classification task in which interclass differences are known *a priori* to determine whether QuAC is able to correctly identify them. To that end, we created a synthetic dataset by simulating individual confocal z-slices through cells with three fluorescent channels that target the cell membrane (green), the nucleus (blue), and a set of punctate objects (red) that represent small organelles or protein aggregates (see Appendix A). We created the dataset to have two axes of variation: the elongation of the cell (round vs. elongated) and the distribution of the punctae (diffuse vs. aggregated), thus resulting in four different classes (Fig. 2a). While these axes perfectly describe the classes, intra-class variation is high enough to make the differences subtle at times. We highlight with a white star some examples in Fig. 2a that would be difficult to tell apart for a human annotator.Figure 2.

**Figure 2.**
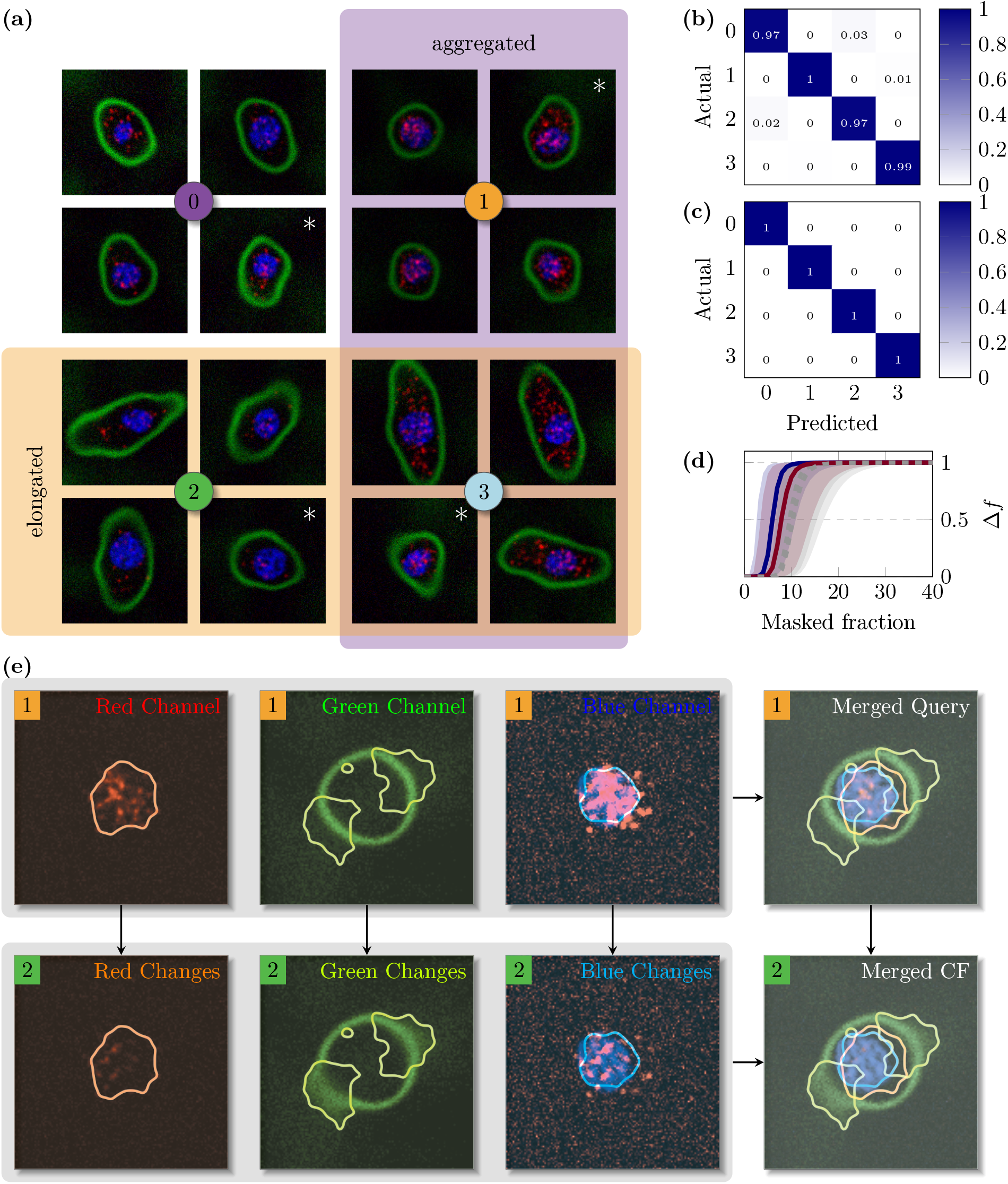
Simulated dataset with known features. **(a)** This data set was created as a sanity check for QuAC. Two features distinguish between the classes: elongation of the cell membrane and aggregation of the punctae in the red channel. **(b)** The classifier is able to distinguish between the classes, as shown in the confusion matrix. *(caption continued on next page)(continued)* **(c)** We are able to get a 100% conversion rate regardless of source or target class because we can sample generated images. **(d)** We show the QuAC curves for Discriminative Integrated Gradients (navy blue) and Discriminative DeepLift (burgundy). Both attribution methods are able to explain the classifier’s decisions well with a small area, with the Integrated Gradients being slightly more precise. For completeness, we also show the non-discriminative curves in gray. **(e)** Here we show an example conversion, from source 1 to target 2. The top row corresponds to the query image, and the bottom row is the counterfactual. In the red channel, we see that the punctae get dimmer and less aggregated. In the green channel, the cell membrane gets more elongated - this is done by increasing the fuzziness of the membrane at the edges. In the blue channel, we also get unexpected evidence of cell elongation: the nucleus gets slightly smaller. The features remain visible in the merged images; they all correspond to expected differences for 1 *→*

Despite this, a ResNet18 classifier trained with binary cross-entropy almost perfectly classifies images into the four classes (Fig. 2b). Furthermore, we observe perfect performance of the translation of real query images of one class into generated images of another class (Fig. 2c). The QuAC curves show that several discriminative attribution methods (Section 3.2.2) are able to identify localized class-relevant features (Fig. 2d). Our discriminative variant of the Integrated Gradients method achieves the highest QuAC score; the counterfactual images are generated by modifying no more than 10% of the original query image.

As in Section 1, the chosen attribution is turned into a binary mask by thresholding and smoothing. The threshold is chosen to obtain the smallest possible mask, with the largest effect on the classifier’s decision. Here, the mask has three channels to match the three channels in the images. In the example in (Fig. 2e), we translate from class 1 to class 2; we show these channels separately, and outline the regions of change. The expected changes are a decrease in the aggregation of the punctae, which is visible in the red channel, and an increase in the cell elongation, which is visible in the green channel. Interestingly, the cell elongation is also hinted at in the blue channel, where the nucleus slightly reduced in size, hinting at a movement out of frame. Indeed, in a larger cell, the nucleus would not necessarily be centered in the central z-slice.

In Fig. 3, we show the most informative (*i*.*e*. highest QuAC score) query-counterfactual pairs for every translation to and from class 0. In Fig. 3a, the outlines show that the changes are limited to the red channel. The punctae in this channel become brighter and more aggregated in the counterfactual than they were in the query. Conversely, in the opposite direction (Fig. 3b), the punctae become more diffuse. In Fig. 3b, we also see a slight tightening of the cell membrane, which is not visible in the other direction.Figure 3.

**Figure 3.**
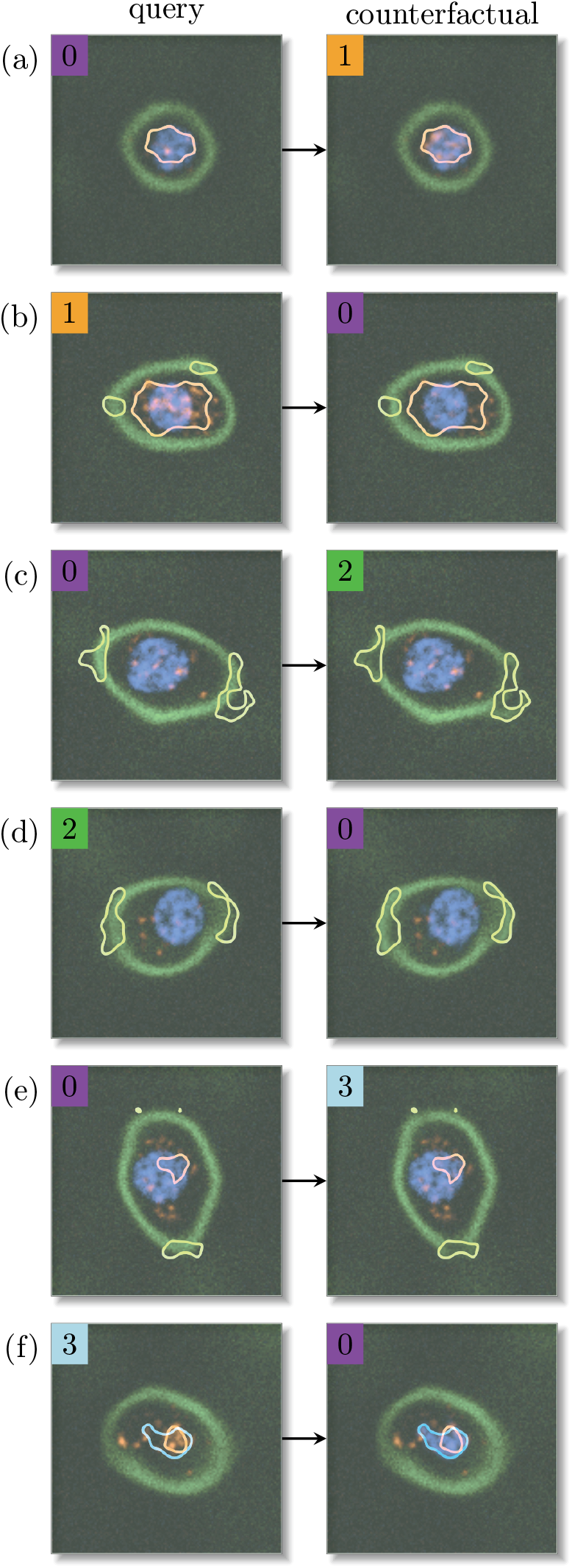
Examples of counterfactual explanations for the Fictus dataset. Here we show the merged images for every direction to and from class 0. **(a)** 0 → 1: there is an increased brightness of the punctae in the red channel, as expected. **(b)** 1 → 0: the punctae in the red channel are conversely dimmed, as expected. There is also a slight change in the green channel, which is unexpected. **(c)** 0 → 2: the cell membrane is increased, as expected, by fuzzing the green channel at the membrane edges, or moving them. **(d)** 2 → 0: the cell membrane is made denser on the edges, signalling a decrease of the cell elongation, as expected. **(e)** 0 → 3: the cell membrane is increased by fuzzing, and the brightness of the punctae is increased. Both of these features are expected. **(f)** 3 → 0: the brightness of the punctae is decreased, as expected. The cell elongation is not affected through the membrane (green) channel, but rather through the nucleus (blue) channel. Specifically, the nucleus is brought into frame, as it would be if the cell was less elongated. This is an expected feature (elongation), appearing in an unintuitive manner.

The changes in both Fig. 3c and Fig. 3d are limited to the green channel. From 0 → 2, we see a slight widening of the cell membrane along the long axis of the cell: the cell is becoming more elongated. In the opposite direction(Fig. 3d), the membrane becomes denser at the edges, suggesting a rounder shape.

Fig. 3e shows changes in both the green and red channels. In the red channel, large, bright agglomerations of punctae are being added in a very limited area of the cell. Simultaneously, in the green channel, the edge of the membrane at the long axis is being pushed slightly outwards and made fuzzier, suggesting an elongation. From 3 → 0, we again see changes in the red channel, this time with existing punctae being dimmed. While there is no change in the green channel, we see a nucleus appear in the blue channel. This is akin to the example of Fig. 2e: the nucleus being pushed into the central frame suggests a smaller, rounder cell.

Combining these observations suggest, we could create a hypothesis that class 0 is defined by cells that have a round shape and more diffuse punctae. These two features are indeed what we used to define the class (Fig. 2a, Appendix A). The QuAC method allows us to retrieve these features from observation of differences between query and counterfactual. We show the query-counterfactual pairs describing the other classes in the appendix (Appendix A).

### 2.2 Electron Microscopy Images of *D. melanogaster* Synapses

Here we consider a dataset of synapses imaged with electron microscopy (EM). Each synapse is labelled with one of six neurotransmitters which is known to be emitted by the neuron to which that synapse belongs. Fig. 4a shows four examples of each type of synapse in our dataset. EM data is rich in detail, and it is not evident by looking at these images what the synapses within any given class have in common that discriminates them from the synapses in other classes. In other words, this classification task is one that is not yet possible for humans, but can be done with high accuracy by a neural network [2]. Indeed, even on 2D slices of EM centered on the synapse, we obtain very good classification results (Fig. 4b). Furthermore, the translation of query images into images of another class using our method generates correctly classified images (Fig. 4c). As such, we can use QuAC to answer the question of which features the classifier uses to make those accurate predictions.

**Figure 4.**
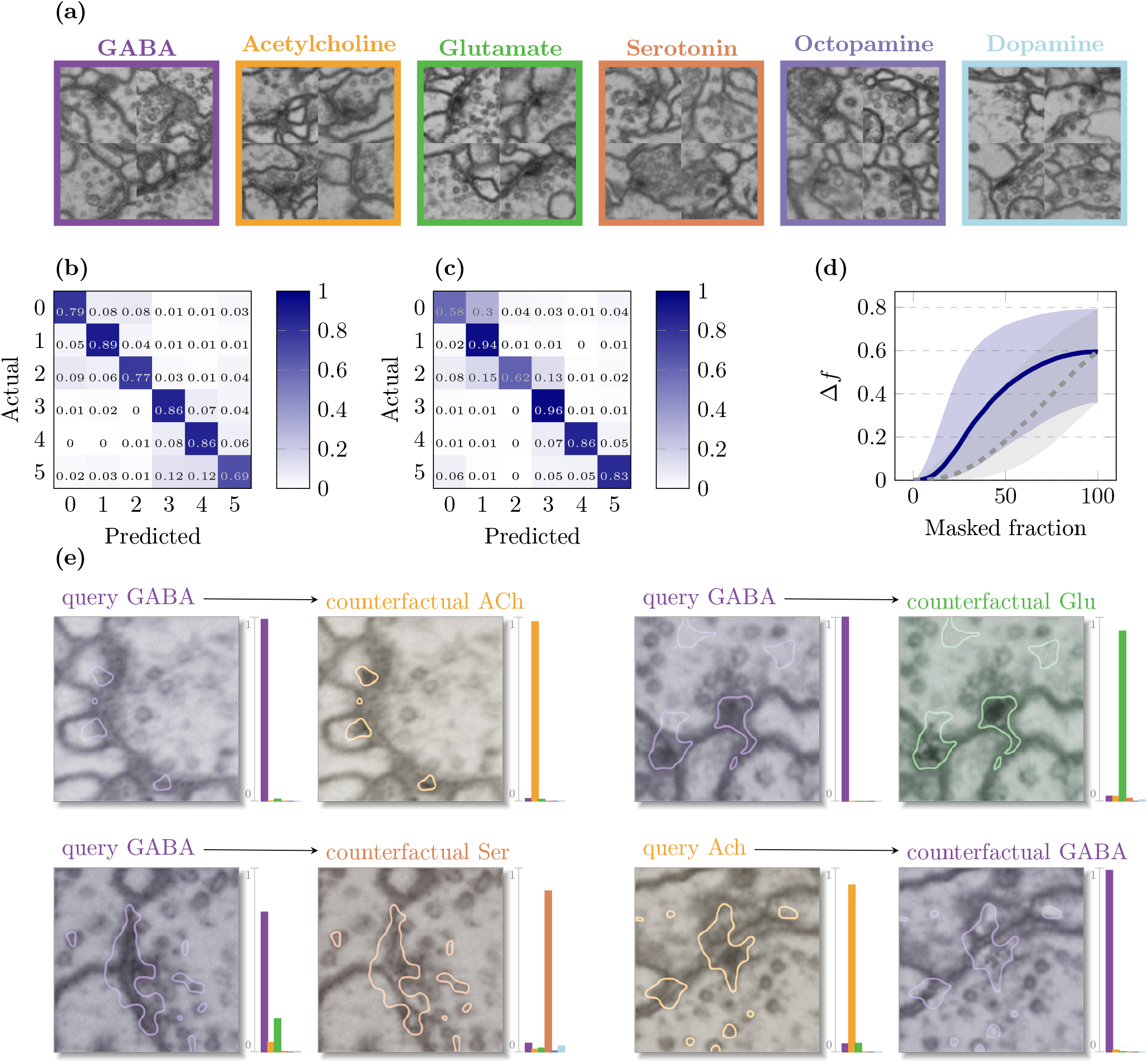
Recognizing neurotransmitters of synapses in electron microscopy images. **(a)** A set of example images for each class of synapse. The classes correspond to neurotransmitters emitted by the synapse. Each synapse is essentially unique, and there is little that allows one to distinguish between classes at first glance. **(b)** Nevertheless, this dataset is highly classifiable as shown in [2], and quickly recapitulated for the 2d network here. **(c)** The classifier has, clearly, learned to distinguish features between these six classes. Training a StarGAN is possible, and leads to pretty high conversion rates. **(d)** In this complex dataset, using a discriminative attribution (blue, solid line) is strongly preferable over the vanilla version (gray, dotted line). Over the full validation dataset (including some bad conversions!) we can explain the differences between classes using less than half of the pixels in an image most of the time. **(e)** We once again show examples with highest QuAC scores for a set of source-target pairs. The focus is generally highly reduced into a specific feature or two of the image, which are very amenable to annotation.

Although an early version of this method was already shown to provide verifiable biological differences between classes [2], we find here several *new* hypotheses. A set of explanations are shown in Fig. 4e, along with the classifier’s output for both the query and the counterfactual image. We see that the focus is on a limited set of features for each synapse. Conversion from GABA to Acetylcholine is driven by a darkening of the post-synaptic densities. These are conversely brightened in the other direction, combined with a removal of the T-Bar, a tightening of the cleft (previously verified), and a removal of some vesicles. GABA-ergic synapses are also characterized by a thinner cleft, lighter T-bar, and fewer vesicles in comparison to glutamatergic and serotinergic synapses (examples 2 and 3), suggesting that these features register as distinctly GABA-ergic to the classifier. Put together, these observations suggest that GABA-ergic synapses are described by a thin cleft, a small or non-existent T-bar, and fewer vesicles and post-synaptic densities. This description could be refined for GABA and created for neurotransmitter types, with the annotation of more examples in the future.

### 2.3 Kenyon Cell classification

An interesting feature of QuAC is that it can be used to interrogate the decisions of a classifier *even in cases where the classifier is wrong*. While the synapse classifier is generally accurate, there are situations in which it is consistently wrong. One such case is the classification of Kenyon Cells in the *D. melanogaster* mushroom body, which are consistently classified as dopaminergic. In Fig. 5a, we show a sample of 500 (of a total of 5177) Kenyon cells from FAFB-FlyWire [3, 4], colored by the classifier’s prediction, where dopaminergic cells are shown in light blue. Importantly, these cells are actually cholinergic [5].

**Figure 5.**
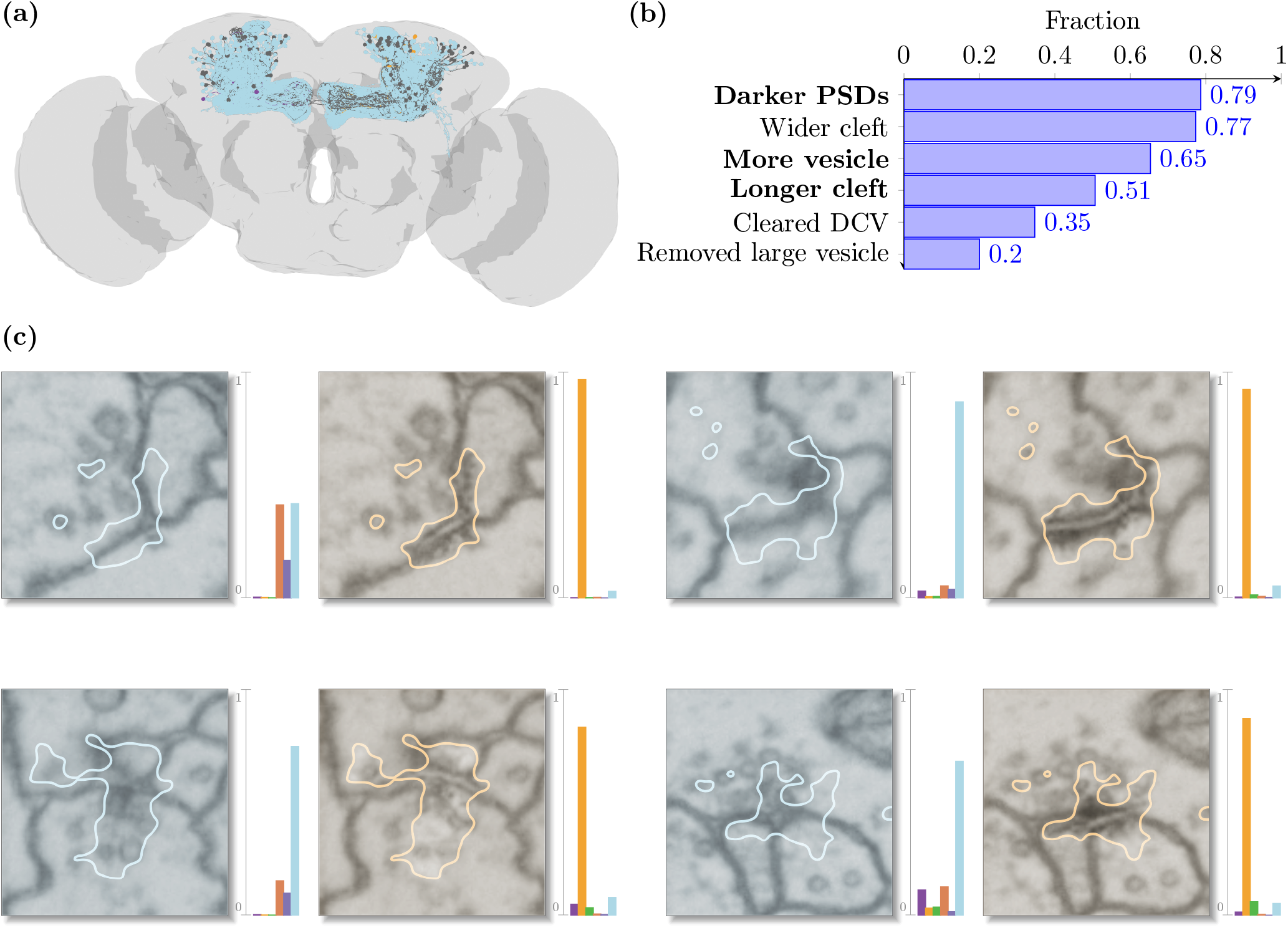
Understanding the misclassification of Kenyon cells. **(a)** A sample of Kenyon cells placed in the brain, colored by how they are classified by the 2D VGG model. The large majority of these cells are classified as Dopaminergic (light blue) although they are Cholinergic. Also visible are some cells classified as GABA-ergic (purple), some correctly classified as Cholinergic (gold), and some cells with too low a confidence for a classification, noted as “unknown” (gray). **(b)** A quantificaton of the features found when turning the Kenyon cell synapses from Dopaminergic to Cholinergic and attributing with QuAC. Bolded features are were already found in [2] to describe the difference between Acetylcholine and Dopamine. **(c)** Example images and their counterfactuals, showing the features described in **(b)**.

**Figure 6.**
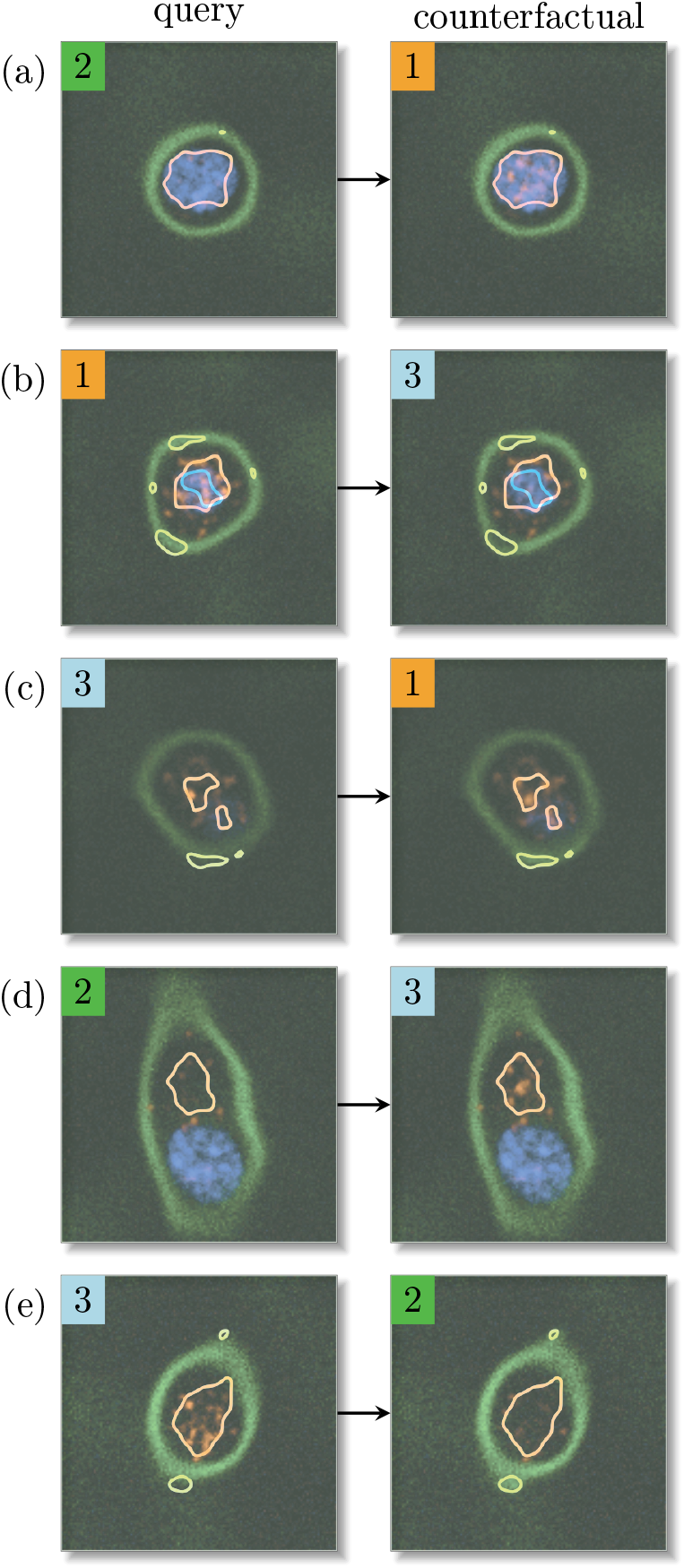
Remaining examples of counterfactual explanations for the *Fictus* dataset.

**Figure 7.**
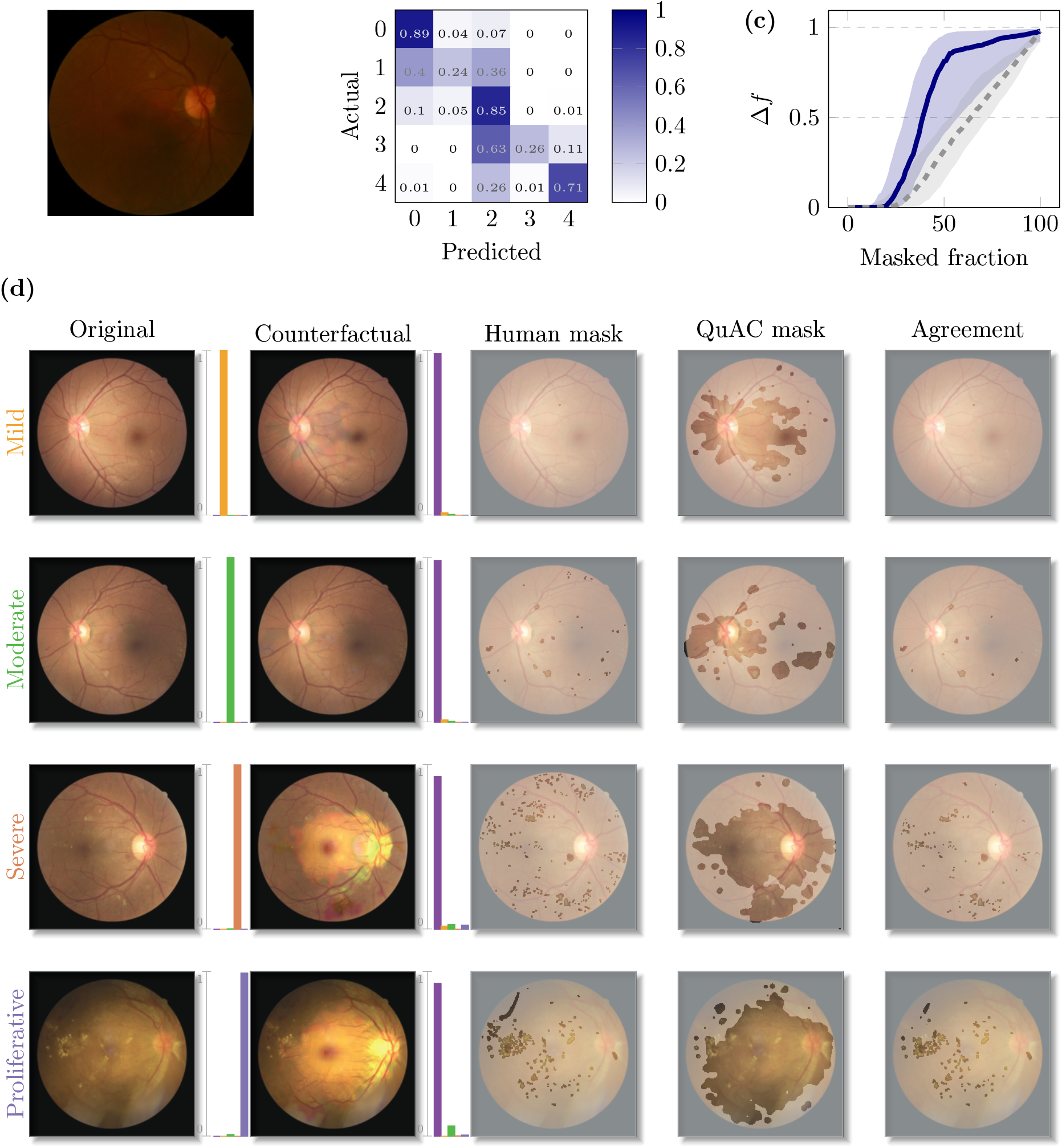
Comparing human and QuAC explanations for retinal images. **(a)** An example of a healthy retina. **(b)** Confusion matrix for the source images for the best classifier. **(c)** QuAC scores for the case study, showing Discriminative Integrated Gradients, as well as a vanilla version for comparison. **(d)** Examples for each level of disease severity, showing the original image, the counterfactual, and the explanation from both human annotation and from QuAC. We highlight the regions of agreement between the two explanations.

The misclassification is consistent across multiple classifiers, on various connectome datasets, in both 2D and 3D (see Table 2). This suggests that the structural features present in Kenyon Cells do not match those of other cholinergic neurons in the dataset, but rather more closely resemble those of dopaminergic neurons. We find that we can verify this hypothesis by using QuAC.

**Table 1:** The two axes of variance put into the *Fictus* dataset. The elongation ratio is the ratio of the large axis to the small axes of the membrane. The 2D slice chosen for this work is not necessarily along the long axis. The number of clusters is the expected number of clusters in the red channel. The actual number is sampled for each cell from a normal distribution with a standard deviation of 1.

| Class | Elongation ratio | Number of clusters |
| --- | --- | --- |
| 0 | 1.5 | 50 |
| 1 | 1.5 | 200 |
| 2 | 2.5 | 50 |
| 3 | 2.5 | 200 |

**Table 2:**
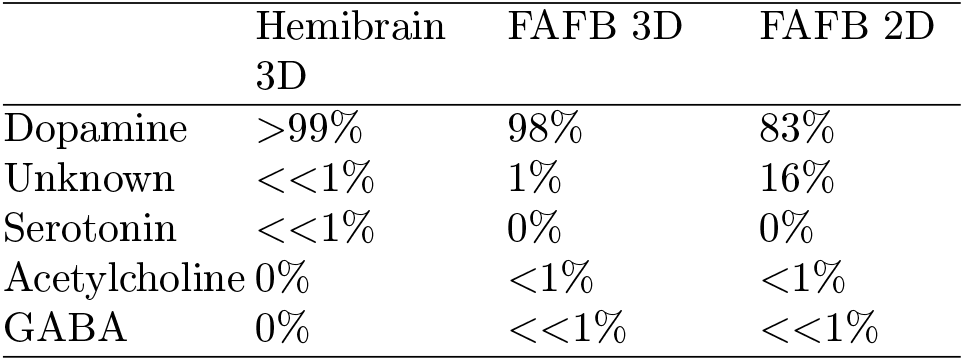
Kenyon cells classification results.

|  | Hemibrain 3D | FAFB 3D | FAFB 2D |
| --- | --- | --- | --- |
| Dopamine | >99% | 98% | 83% |
| Unknown | <<1% | 1% | 16% |
| Serotonin | <<1% | 0% | 0% |
| Acetylcholine | 0% | <1% | <1% |
| GABA | 0% | <<1% | <<1% |

To do this, we translated a set of 100 Kenyon cell synapses from dopamine to acetylcholine. Using these images, we first compiled a set of discriminative features between the (mis-classified) query image and the (correctly classified) counterfactual. A complete list of selected features, as well as a rubric for when to annotate them, can be found in B.2. We then manually assigned these features to each of the hundred synapses to quantify prevalence. Where there was uncertainty in the manual annotation, we removed the synapse from analysis, leading to *n* = 75.

We recapitulate the top 6 features, each found in at least 20% of the annotated synapses, in Fig. 5b. Among these, we find three that were known differences between acetylcholine and dopamine (from [2]), highlighted in bold. Three of the distinguishing features were new, including a widening of the cleft which was highly prevalent among the annotated synapses. A similar manual annotation of *real* dopaminergic synapses would determine whether these new features are unique to Kenyon Cells.

In Fig. 5c, we show examples of how these features manifest in the images. As was the case in Section 2.2, the features are subtle and would be difficult to detect without the paired counterfactual.

## 3 Method

### 3.1 Related Work

Existing post-hoc explainability methods can broadly be grouped into two categories: gradient-based methods and latent-space methods.

Gradient-based attribution methods use back-propagation of gradients through the network to determine either the input-level pixels [6, 7, 8, 9, 10, 11, 12] or the internal activations [13, 14, 15, 16, 17] that have the strongest effect on the classifier output and are therefore expected to be the most important. The effective relevance of the regions found by these methods is difficult to quantify and verify: While it is possible to quantify how the degradation of these regions affects the classification [18], it is not possible to tell whether the drop in performance is due to the importance of the degraded region, or simply due to the image being degraded in the first place. Furthermore, attributions from gradient-based methods are difficult to interpret as they only highlight presumably relevant regions, but do not answer why a region is considered important. Such methods might, for example, consistently highlight the nucleus as being important for the classification of cell’s state, but will not reveal what feature of the nucleus (*e*.*g*., intensity, texture, or shape) is discriminative. While it is possible to perturb an image in the direction of the gradients to visualize changes [19], such methods require robust classifiers to avoid obtaining adversarial images that merely exploit vulnerabilities in the neural network but are unlikely to represent indistribution samples. QuAC extends gradient-based methods as in [1] by making them *discriminative* and providing a *quantitative* measure of the quality of the highlighted regions.

In contrast, latent-space methods aim to identify high-level features (*e*.*g*., cell size, shape, and intensity variations) of a dataset. Latent-space methods are trained, often in a self-supervised manner, to create representations of the data that are smaller, more easily navigable, and empirically more interpretable than the input images [20, 21]. Latent representations can also capture more complex features than standard dimensionality reduction methods (*e*.*g*., PCA). Indeed, latent-space analysis of variational auto-encoders and transformers have been shown to provide valuable biological explanations [22, 23, 24]. However, these methods are generally not designed to work hand-in-hand with a pre-trained classifier. Instead, they provide a representation of the data that is unbiased, and therefore does not differentiate between class-relevant and class-irrelevant features. There are notable exceptions [25, 26], where pretrained classifiers are used in conjunction with latent space methods to identify mostly independent latent feature dimensions that affect classification. Similarly, QuAC includes a latent space (see **style**) that is specifically trained to contain just class-relevant features, combined with a powerful image generation model to handle all the class-irrelevant features.

### 3.2 QuAC

Our method presents an alternative to gradient-based and latent-space explainability methods by generating plausible and interpretable counterfactuals. Given an image of one class, the counterfactuals for that image show which parts of the image have to be changed, and in which way, such that the image will be classified as belonging to another class. We first train a separate neural network (a variant of the StarGAN [27], see Section 3) to convert images between different classes. During training of the converter, a discriminator ensures that the generated images are representative of the target class, *i*.*e*., that they are in-distribution samples. Furthermore, the converter is incentivized to make minimal changes to the image, such that class-invariant features like the location or orientation of a structure of interest are retained, but class-variant features are changed. Importantly, the converter is trained independently of the classifier to avoid obtaining adversarial samples that exploit classifier weaknesses. We next find an *attribution map* that assigns a real-valued score to each pixel, representing how important the change of that pixel (between the real and generated image) was in changing the classifier’s prediction. We then threshold the attribution map at several levels to obtain a series of binary masks, each highlighting increasingly larger regions of the most important changes (see Fig. 1b). We can copy the masked content of the generated image into the original real image, and classify the result. Importantly, we can quantify the amount by which this region changes the classifier’s prediction as compared with the original image. We can then search for the smallest mask that highlights the most important differences. We obtain the counterfactual using only this minimal masked region. The counterfactual thus differs from the original real image only in the masked region, but is classified differently.

#### 3.2.1 Image generation with StarGAN

The basis for this pipeline is the targeted, conditional generation of images using a StarGANv2 model [27]. By targeted, we mean that the images are generated to have a specific class. The StarGAN’s *generator G* takes as input the image *x* and a *style*, which is a representation of the target class *y*_*t*_. It then returns a realistic, generated image *x*_*g*_ that looks like the input except for some key features. The trained classifier *f* should classify *x*_*g*_ as the target class: *argmax f* (*x*_*g*_) = *y*_*t*_.

Two networks are used to create the style representation: the *mapping network* and the *style encoder*.

The *Mapping network M* takes as input a randomly sampled latent code. It has as many outputs as there are classes, and each is a style vector for a different class. Only the style for the target class *y*_*t*_ is passed to the generator.

The *Style encoder E* takes as input a real image and creates a style. Specifically, the style should be a representation of the class of the image. We created two different versions of the style encoder. The default encoder has as many outputs as there are classes, like the mapping network. Similarly, only the style vector for the target class *y*_*t*_ is passed to the generator. We also create a single-output version of the style encoder, which is not explicitly given the target class. In order to encode class information, the single-output style encoder needs to learn to recognize the class directly from its input.

As the StarGAN is an adversarial model, a *discriminator D* is trained concurrently with the generator. The purpose of the discriminator is to recognize real from fake images. Following [27], the discriminator also has one output for each class - only the output corresponding to *y*_*t*_ is used at any given time.

There are three losses and two regularizations used to train the StarGAN model. The generator and discriminator compete with an adversarial loss, for which we used the LSGAN loss. The generator is also trained to keep the generated image as close as possible to the input image with a cycle consistency loss. Given an image *x* and a target style 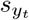 the cycle consistency loss ensures that we can recover *x* after it has been translated to the target style and back.

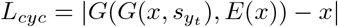

Similarly, there is a style consistency loss:

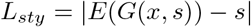

In order to encourage diversity in the effects of styles on the generated images, a style diversification regularization is used. This is a L1 distance between two output images that have the same content image as a source, but different styles. Maximizing this loss encourages the generator to use different styles for different images, but it can create unwanted artifacts which we discuss below. Finally, an R1 regularization is used on the discriminator.

Throughout training we keep an exponential moving average the generator, mapping network, and style encoder. At every step *i*, this weights *w*_*ema*_ of the exponential moving average models are updated as an interpolation between the previous EMA weights and the weights of the training model *w* as follows:

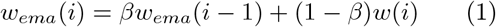

We use *β* = 0.999. The EMA models are used to generate images during inference. This makes the outputs more stable and realistic.

One of the advantages of the StarGAN over the CycleGAN used in a previous iteration of this method [1] is the ability to sample generated images. We define the *translation rate* as the fraction of generated images correctly classified by the classifier during inference. However, for each input image we can sample multiple generated images, each with a different style. We do this, generating at most 256 images per input image, and then select the one with the highest target class probability. We define the *conversion rate* as the fraction of generated images sampled in this way that are correctly classified by the classifier during inference. In a deterministic model such as the CycleGAN, the translation rate and conversion rate are the same. In the StarGAN, this sampling allows us to increase the conversion rate simply by sampling more generated images. This leads to a much larger number of samples that can be used for counterfactual generation and attribution.

#### 3.2.2 Discriminative Attribution

We determine the most important differences between an image and its counterfactual by using discriminative attribution. A discriminative attribution method [1] is built upon a classical attribution method, but uses the counterfactual image as a comparative. For methods that assume a baseline [11, 8], the counterfactual image can be used as a baseline. Other methods were modified according to [1]. These methods return a heatmap of how much each pixel contributes to the classifier’s decision between the source class, which is the class of the original image, and the target class, which is that of the counterfactual. We process the attributions so that they represent contiguous regions, rather than independent pixels. We can apply a set of attributions on the same data and counterfactuals, as we can use the QuAC score to choose the most informative one (see Section 3.2.3). For completeness, we also show the results for non-discriminative versions of the attribution methods, although these were consistently shown to be worse in [1].

Importantly, because we are able to evaluate the attributions, we are not limited to choosing a single method for our whole dataset. In this work, we used Integrated Gradients and DeepLIFT, both the single-image and the discriminative versions, and choose the best attribution for each image. We limited ourselves to these two methods because of computational constraints, but any method that returns an attribution map of pixel contributions could be considered here.

#### 3.2.3 Mask, counterfactual, and evaluation

We find the mask based on a direct quantification of the effect of that region on the classifier. We can obtain a series of binary masks from a single attribution map by varying the threshold of importance that we wish to evaluate. We can then use these binary masks to create hybrid images: the hybrid image contains some pixels from the input image (masked-out region) and others from the counterfactual (masked-in region). Importantly, this hybrid image is still indistribution and can be reliably classified. Qualitatively, if the hybrid is of class A then our mask does not cover all the discriminative features.

The attribution is thresholded at different levels to create a set of binary masks *m*. Each binary mask covers an increasingly large region of the image, which the smallest mask covering only the most important regions according to the attribution method. We then create a hybrid image *x*_*h*_ for each mask: where the mask is 0 we put the pixels from the original image, and where the mask is 1 we put the pixels from the counterfactual image.

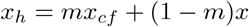

The hybrid is an image that we can directly classify, and the classification of the hybrid lets us determine how good of a mask we have made. If the mask is too small, the hybrid should still be classified as the source class. As soon as it is large enough to capture the discriminative features, the hybrid’s classification should change to the target class.

We can directly quantify the size of the masks to its effect on the classifier’s decision by looking at *f* (*x*_*h*_). In particular, we can look at the change in the classifier’s output as a function of the mask size. We can then calculate the QuAC score as the area under this curve. The quality of our explanation can be also be *quantitatively* scored by comparing the change in classification to the normalized size of the binary mask. We obtain the QuAC score as the area under this curve; this score allows us both to choose an attribution map and to rank the samples by how informative their features are about the entire class. We choose as the optimal mask – and consequently the optimal hybrid – the smallest mask with the highest effect on the classifier output. The optimal hybrid is our counterfactual image, which shows the minimal changes needed to change the classifier’s decision. This mask and counterfactual are then used to select and describe the features of interest.

#### 3.2.4 Code availability

The code for QuAC is available here. The code for the various experiments can be found here.

## 4 Discussion

Neural networks are powerful tools for classification in biology, but they are often seen as black boxes. We developed a method that allows us to reverse engineer these black boxes and visualize the visual features they use to make decisions. QuAC combines counterfactual generation with discriminative attribution to create a setup that allows for relatively easy quantification of visual features that discriminate between classes.

The quality of the discriminative attribution, and therefore the usefulness to find new insights into datasets, depends on the accuracy of the classifier under investigation. The highlighted regions in paired images are most informative if the classifier achieves a sufficiently accurate separation of classes. Consequently, a mediocre classifier will lead to attributions that still explain the decision boundary of the classifier but are likely not representative of the data distribution. We give an example of this in Appendix C. Conversely, good classifiers will lead to the discovery of features that are easy to interpret and represent actual differences between two distributions of images. Furthermore, as we showed on synthetic data, QuAC can fully recover all relevant inter-class features under ideal conditions.

Furthermore, the cornerstone of the method is the ability to generate counterfactual images. As such, the quality of the image generation is crucial. A poor generator will not be able to correctly convert images from one class to another, leading to a limited number of counterfactual images. Although annotation is still useful in these cases, the features obtained from limited data will not be comprehensive enough to explain the classifier’s accuracy [2]. Indeed, the features used by neural networks are many and sometimes complex; they must be verified on a representative sample of the data.

In this work, we used a StarGANv2 because it gives us the ability to sample from generated images — this allows us to search for correct conversions. In some cases where there is a difficult conversion, we can choose to increase compute in order to obtain more examples. In other cases, where the conversion is easy, one might consider comparing various explanations for the same original data point. Alternative generators, such as diffusion models, could be used for image generation in the future [28].

In QuAC, the explanation we provide is **quantified**. Previous attempts to benchmark interpretability have been mostly based on degradation of the important parts of images [18]. While useful, these methods do not separate between the degradation effect from finding the features of importance and the effect of creating out-of-distribution images through degradation. We avoid this balancing act by generating images that are within distribution at every step of the way. The only limitation to our quantification is therefore the quality of the classifier itself.

Furthermore, as we show in the Section 2.3, it is also possible to use QuAC to interrogate misclassifications. This is particularly interesting in cases, such as that of the synapses, where the classes that are available for supervised classification do not capture the full heterogeneity expected in the data. In biological data, categorization can generally occur at one level of granularity. This this case, while Kenyon Cells are cholinergic, it may be that they differ enough from other cholinergic neurons to be considered a separate class. Is contrast to the usual solution of ignoring mis-classified samples for analyses, we show that a generally good classifier making misclassifications can be an impetus to dig deeper and understand the biological particularities of the data.

This paper is by no means an exhaustive description the types of data that QuAC can be applied to. Though all the examples given here are image data, we expect that the very same steps can be applied to differently-dimensioned data such as RNAseq or calcium recordings of neurons. We also expect this method to be applicable to time-series data with minimal modifications — most of these modifications to the synthetic data generation process. Furthermore, there are several simple but potentially powerful ways to dig deeper into the method’s outputs for domain-specific purposes. These include looking at multiple counterfactual explanations, exploring the separations of features in the style space, or injecting additional metadata information into the generator.

There are, however, some cases where using QuAC would be inadvisable. First, the final step of the QuAC pipeline remains human annotation. There is never a guarantee that the differences between classes are human-interpretable, and in some cases they may be so subtle that they are not distinguishable by eye. In these cases, the method will not be able to provide any insights.

Furthermore, generative models such as StarGAN are trained to recapitulate the data distribution as well as possible. This means that any signal present in the original data must be present in the generated data as well. This includes modelling confounders in the data — elements that are correlated with the class labels, but that have no biological relation to them. Though some confounders (*e*.*g*. uneven lighting, different magnifications, etc.) can be accounted for and removed through data augmentation or cleaning, there are often unexpected correlations that need to be accounted for, rather, during the data acquisition.

Overall, we expect QuAC to become a tool for probing both new and existing classifiers and generating a wealth of human-interpretable and verifiable hypotheses. We provide the source code, as well as tutorials for testing QuAC on new data. We also provide a simple annotation tools to make use of the QuAC outputs for gaining new biological insights. We expect further improvements to the pipeline to be quick and easy to prototype within this framework. Finally, we hope that the availability of explainability tools such as QuAC will promote new data acquisition practices for better integration with explainable deep learning.

## 5 Acknowledgements

We thank Caroline Malin-Mayor for help in method refinement, as well as help editing and reviewing this manuscript. We thank Manan Lalit, Alexander Hillsley, and Larissa Heinrich for their help in model refinement, and Hugo Senetaire, Alán Fernando Muñoz González, and Hugo Hakem for implementation details and debugging. We acknowledge the CVML journal club at Janelia for their feedback. This work was supported by the Howard Hughes Medical Institute.

## Appendix A Details on the *Fictus* dataset

All the cells in the *Fictus* dataset were created in 3D within a bounding box of size 128 on each side. The code to recreate the dataset can be found in this repository.

We first create the membrane in the green channel. First, we define a certain elongation ratio. The membrane is then created as a 3D ellipsoid, where two of the radii are equal (small axes), and one is larger by the elongation ratio (large axis). The cell is then randomly rotated in 3D space. We then add a jitter to the membrane, so that it is not perfectly elliptic. Finally, we define a width and a level of “fuzziness” for the membrane, which are applied as a sigmoid function to the distance from the membrane center.

Next we create the nucleus in the blue channel. The nucleus is defined as a sphere with a radius equal to 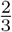 of the small axes of the membrane. The nucleus is place at the point with the highest distance from the membrane. We add jitter to the nucleus membrane so that it is not perfectly spherical. Then, we create a chromatin structure in the nucleus structure by adding gaussian noise within the segmented nucleus, then applying a gaussian filter on top to smooth it. The brightness in the nucleus is proportional to the amount of chromatin within.

Then, we create the punctate structures in the red channel. We randomly sample a number of punctae clusters, then randomly sample their location with an expected distance from the membrane: these will represent clusters of a protein. We then sample a number of individual points to go within each clusters, and randomly assign them to one of the clusters. The brightness of a cluster is proportional to the number of points within it. We apply some gaussian smoothing on top of the punctae to simulate the growth of the protein cluster.

Finally, we simulate the imaging process. We simulate a point spread function that is anisotropic, with a larger spread in the z direction. We then add gaussian noise to each channel independently. In the membrane channel, we also add low-frequency noise to simulate out-of-focus objects.

There are 25000 cells per class in training, and 2500 in the test set.

## Appendix B Kenyon Cells

### B.1 Classification results

The Kenyon cells are consistently mis-classified by multiple classifiers, on various connectome datasets, in both 2D and 3D. The classification results are shown in Table 2.

### B.2 Annotation

Below is the rubric used to annotate features in Fig. 5.

- **Wider cleft**: The cleft got brighter/increase in the direction of the synapse (pre-to-post). Generally, this means that there was just a line, and then a visible cleft was created.
- **Longer cleft** or, equivalently **More postsynaptic partners**: The cleft size increased perpendicular to the synapse (so along, rather than across). This means the visible cleft is now touching more post-synaptic partners.
- **Darker PSDs** or, equivalently **Denser PSDs**: The post synaptic densities either appeared, increased in size, or became less diffuse
- **More vesicle** meaning **Darker spots/added vesicles/larger vesicles**: Either barely visible spots came into focus/darkened or vesicles were created from nothing. This annotation was only considered on the pre-synaptic side.
- **Cleared DCV**: A larger dark vesicle was turned lighter, emptied, or burst/split. This annotation is not used for smaller vesicles.
- **Removed large vesicle**: A larger (clear) vesicle was removed/reduced in size. If reduction in size, often combined with **More vesicle**
- **Added spots**: Spot-like (smaller than vesicles) structures appear, generally around the T-bar area or near the cleft. These spots can appear on the pre- or post-synaptic side.
- **Spots removed**: Spot-like structures disappear from the cleft area, or are transformed into PSDs/T-Bar.
- **T-Bar modified**: The T-Bar was either removed, or made lighter.
- **T-Bar added**: A T-Bar was added where it was not visible before, or increased visibly in size.

## Appendix C Diabetic Retinopathy

### C.1 Introduction

On retinal fundus images we show that features used by humans and features used by the classifier can surprisingly differ.

### C.2 Results

Diagnosing diabetic retinopathy is a popular benchmark task for classification by deep learning models, thanks in part to the availability of large datasets. Experts score the severity of the disease on a scale from 0 to 4, with 0 being healthy and 4 being proliferative diabetic retinopathy. There are several features that are known to be indicative of the severity of the disease, such as microaneurysms, hemorrhages, and exudates. These can also be manually annotated by domain experts, although this is a time-consuming process. We ask, then, whether a classifier trained on this task uses the same features as humans to make its decisions.

We use data from [29], without considering any of the ungradable samples, and replicate their methods to train a ResNet18 classifier. The classifier reaches a balanced accuracy of 59% which is competitive with that in [29] (57% on the same model, ignoring the ungradable accuracy). The majority of the confusion comes from differentiating neighboring severity levels (see Fig. 7b). The same data is then used to train the StarGANv2. Neither the classifier training nor the StarGAN training includes any information about lesion segmentations; these segmentations are only used to compare to the masks found by the attribution method. We discuss the correlation between lesions and predicted severity in more detail in the supplementary material.

For testing, we use the data with lesion segmentations from [29]. The samples in this testing set are not graded, but all of them have at least one type of lesion segmented. We therefore assume that none of the samples are healthy, and use the classifier to predict the severity of the disease. We then use QuAC to convert the samples to the healthy (No DR) class, to determine what the classifier uses to classify disease against no disease. Using discriminative integrated gradients for attribution (see population scores in Fig. 7c), we generate the counterfactual images.

In figure Fig. 7d, we show one example for each severity level, as well as its corresponding explanations. The counterfactual images are all predicted as healthy by the classifier, but all of them require a large change in the image to do so. Indeed, when we compare the lesion segmentations (third column) with the masks found by the attribution method (fourth column), we see that although the QuAC masks cover the majority of the lesions, they also cover large fractions of the image that are unused by human experts. This is especially true for the samples with higher severity levels. In the supplementary material, we show the precision and recall of the attribution masks compared to the lesion segmentations.

### C.3 Lesion segmentations

Four different kinds of lesions are segmented, and we verify that all of them are correlated with the severity classification given by our trained classifier. The QuAC attribution have a high recall for the lesion segmentations, but a very low precision. This suggests that the classifier is using more information than just the lesion segmentations to make its decision.

